# TRIM5α, TRIM56 and TRIM69 differentially regulate dengue virus 2 infection through selective NS2B3-mediated antagonism

**DOI:** 10.64898/2026.11.16.767120

**Authors:** Ricky Raj Paswan, Luan Letieri Belem Martins, Natacha Merindol, Solena Rossi, Aïcha Aïssatou Sow, Nguyen Ha Nhung, Jyotika Singh, Laurent Chatel-Chaix, Isabel Desgagné-Penix, Lionel Berthoux

## Abstract

Dengue virus (DENV) is a major mosquito-borne human pathogen. Understanding the mechanisms of cellular resistance to the infection is vital to develop novel therapeutic tools. Tripartite motif (TRIM) proteins contribute to antiviral innate immunity through direct restriction of viral replication and regulation of antiviral signaling, but current knowledge on DENV targeting by TRIM proteins remains preliminary. Previously, TRIM56 was found to inhibit DENV, whereas reports on whether TRIM69 is a DENV restriction factor are conflicting. Surprisingly, the broad-spectrum viral restriction factor TRIM5α was proposed to restrict tick-borne but not mosquito-borne flaviviruses. Here, we investigated the effects of TRIM5α, TRIM56, and TRIM69 on DENV serotype 2 (DENV2) infection. Overexpression experiments showed that TRIM5α and TRIM56 restrict DENV2, whereas TRIM69 enhances it. Consistently, CRISPR/Cas9-mediated loss-of-function showed that knocking out TRIM5α or TRIM56 increases infection, while TRIM69 knockout decreases it. Restriction correlated with reduced viral RNA and viral protein levels in TRIM5α- and TRIM56-expressing cells. Virus-free experiments showed that TRIM56 downregulates the DENV2 protease complex NS2B3 levels. Reciprocally, DENV2 NS2B3 reduced TRIM56 and TRIM69 abundance independently of protease activity. RING domain-mutated TRIM56 retained partial antiviral activity and the ability to reduce NS2B3 levels, and TRIM56-NS2B3 reciprocal downmodulation effects were not counteracted by proteasomal or lysosomal inhibition. Together, these findings establish TRIM5α as a DENV2 restriction factor, reveal TRIM69 as a positive factor in DENV2 infection, suggest that TRIM5α and TRIM56 inhibit DENV2 through distinct mechanisms, and hint at proteasome-independent TRIM56-NS2B3 reciprocal downmodulation.

**IMPORTANCE:** DENV is a growing global health concern, with transmissions expanding into new geographic regions. No antiviral drug treatment has been approved for DENV, despite it being the most prevalent mosquito-borne viral infection worldwide. This highlights the need to better understand host factors regulating DENV. Many TRIMs are key components of the intrinsic antiviral defense, but their roles during DENV infection remain poorly understood. Here, we report on the characterization of novel antiviral and pro-viral effects of cytoplasmic human TRIMs on DENV2. We further show that the DENV protease complex NS2B3 decreases TRIM56 and TRIM69 protein levels through a mechanism that remains undetermined but suggests the possibility of a viral counter-measure strategy. Together, these findings demonstrate distinct roles for TRIM proteins in controlling DENV2 and hint at viral resistance mechanisms to TRIM-mediated DENV restriction.

## INTRODUCTION

Dengue is a major mosquito-borne human viral disease, and its prevalence has increased markedly over the past several decades. More than half of the world’s population is currently considered at risk of dengue infection, with the World Health Organization estimating 100 to 400 million cases per year (1). The geographic range of dengue has continued to widen along with the distribution of its principal vectors, *Aedes aegypti* and *Aedes albopictus* (2). Two vaccines have been approved, i.e. Dengvaxia and Qdenga; but their usage is limited by factors such as prior dengue exposure, age, local dengue disease incidence, and virus serotype (3). No antiviral drug is approved for treating dengue, and case management remains supportive (4). Dengue virus (DENV), a member of the *Orthoflavivirus* genus, from the *Flaviviridae* family, comprises four antigenically distinct serotypes (DENV1-4) (2). DENV is an enveloped, positive-sense single-stranded RNA virus with a genome size of ∼11 kb. The viral genome is translated into a single polyprotein and processed by host and viral proteases into three structural proteins (capsid, prM/M, and envelope), and seven nonstructural proteins (NS1, NS2A, NS2B, NS3, NS4A, NS4B, and NS5) (5–7). The nonstructural proteins play important roles in viral RNA replication, polyprotein processing, virion assembly, and the regulation of host cell responses (7). NS3 contains an N-terminal serine protease domain that requires NS2B as a cofactor for its proteolytic activity. NS2B contributes to the folding and stability of the NS3 protease domain and is responsible for NS3 recruitment from mitochondria and anchoring into endoplasmic reticulum membranes (7–9). The NS3 C-terminal region has RNA triphosphatase, nucleoside triphosphatase, and helicase activities (5, 7). NS5 displays RNA-dependent RNA polymerase and methyltransferase activities essential for viral RNA synthesis and RNA capping (5).

Viral molecules are sensed by cellular pattern recognition receptors, leading to activation of innate antiviral signaling pathways (10, 11). For instance, RIG-I and MDA5 detect replicative forms of the DENV viral RNA and signal through MAVS to induce type I interferon (IFN-I) production (12, 13). IFN-I activates the JAK-STAT pathway, leading to the formation of the ISGF3 complex which translocates to the nucleus and induces the expression of interferon-stimulated genes (ISGs) that establish an antiviral state (14, 15). A subset of these ISGs acts as restriction factors, i.e. host proteins that are effectors of the cell-autonomous antiviral immunity by directly limiting one or more stages of virus replication. In addition to being IFN-I-induced, some restriction factors basal expression levels are sufficient for them to act as intrinsic barriers to infection (10). Restriction factors can target different stages of viral replication and through different mechanisms. They can inhibit entry, genome replication, assembly, or egress and have been identified across many virus families (16). For example, IFITM3 restricts influenza A virus by preventing viral entry into the cytosol, OAS1 restricts SARS-CoV-2 replication through RNase L activation, and tetherin inhibits HIV-1 virion release (17–19). Several restriction factors targeting flaviviruses have been identified. These include IFI6, which interferes with the formation of flaviviral replication organelles, viperin, which inhibits viral RNA synthesis through the formation of the nucleotide analogue ddhCTP, and RyDEN, which binds DENV RNA and decreases viral RNA replication as reviewed in (10).

Among virus restriction factors, one of the largest groups is that of the tripartite motif (TRIM) proteins. More than 70 TRIM proteins are found in humans, and they share a conserved N-terminal RBCC arrangement of a <u>R</u>ING domain, one or two <u>B</u>-boxes and a <u>c</u>oiled-<u>c</u>oil domain, followed by a variable C-terminal region that determines target recognition (20, 21). The RING domain confers E3 ubiquitin ligase activity, whereas the B-box facilitates higher-order multimerization and the coiled-coil regions mediates TRIM-TRIM interactions, leading to their dimerization and oligomerization, which upregulates their E3 ligase activity (22). The C-terminal region most often takes the form of a PRY-SPRY (B30.2) domain, though NHL-repeat and other domains also occur, and this variability has led to TRIMs classification into several subfamilies (11, 23). Many TRIMs are IFN-I-inducible (24), and several have been found to inhibit viruses either by acting directly on viral components or indirectly by regulating innate immune signaling (24). For example, TRIM22 ubiquitinates the influenza A virus nucleoprotein and targets it for proteasomal degradation (25), whereas TRIM52 interacts with the NS2A protein of Japanese encephalitis virus and promotes its K48-linked ubiquitination and proteasome-dependent degradation in a RING domain-dependent manner (26). Several human TRIMs were found to target flaviviruses, such as TRIM22, which ubiquitinates ZIKV NS1 and NS3 in a RING domain-dependent fashion, leading to proteasomal degradation, and was also reported to restrict DENV and YFV (27). More recently, TRIM14 and TRIM21 were proposed to restrict both ZIKV and LGTV, whereas TRIM38 was suggested to restrict only ZIKV (28). In addition, some TRIMs are thought to inhibit flaviviruses and other viruses by upregulating IFN-I-dependent innate signaling pathways (29–31).

TRIM5α is one of the most thoroughly characterized antiviral TRIMs. It was first identified as a restriction factor of HIV-1 infection in cells from Old World monkeys through recognition of the incoming retroviral capsid (32). Its PRY-SPRY domain physically interacts with the assembled retroviral capsid lattice, and hexagonally arrayed TRIM5α dimers engage incoming capsids to accelerate uncoating and target them for proteasomal degradation (20, 32). Interestingly, this restriction activity is coupled with an innate-sensing function. Specifically, TRIM5α-capsid interactions stimulate the RING domain, together with the UBC13-UEV1A E2 complex, to generate K63-linked polyubiquitin chains that activate TAK1 and downstream AP-1/NF-κB signaling. Thus, TRIM5α acts as both a direct restriction factor and a pattern-recognition receptor for the retroviral capsid (33). Interestingly, TRIM5α also restricts evolutionarily divergent viruses, as evidenced recently by the report that it can inhibit DNA viruses from *orthopoxvirus* genus. Specifically, the TRIM5α SPRY domain binds the poxviral capsid protein L3, again coupling restriction by activation of innate signaling (34). Several orthopoxviruses counter this activity through the viral protein C6, which binds the RING domain of TRIM5α and induces its proteasome-dependent degradation (34).

TRIM5α has also been proposed to target flaviviruses, though in a highly virus-specific fashion. Specifically, TRIM5α was reported to restrict several members of the tick-borne encephalitis virus (TBEV) serogroup, namely TBEV, Kyasanur Forest disease virus and Langat virus, by ubiquitinating and degrading the viral NS2B3 complex. Interestingly, the same study reported a lack of restriction for the mosquito-borne flaviviruses tested, including DENV2 strain New Guinea C (NGC), Zika virus (ZIKV), yellow fever virus (YFV), and West Nile virus (WNV), or for the tick-borne Powassan virus (35). Recently, it was reported that a single amino acid change in NS3 was sufficient to make the Powassan virus resistant to TRIM5α restriction, and similar NS3 changes also conferred resistance in other sensitive tick-borne flaviviruses (36), identifying this viral protein as a viral determinant for TRIM5α sensitivity.

Other TRIMs have been reported to restrict DENV and related flaviviruses through different mechanisms. TRIM56 is an antiviral TRIM protein with a C-terminal NHL-like region that has been reported to bind viral RNA and restrict several RNA viruses. TRIM56 restricts YFV and DENV2 in a manner that requires both its RING-dependent E3 ligase activity and the integrity of its C-terminal region, and it impedes viral RNA replication (37). TRIM56 also restricts ZIKV, and its RING and C-terminal regions are required for this activity as well (38). In the latter study, the C-terminal region was shown to bind ZIKV RNA directly, independently of RING activity, while restriction itself still required both domains together. In addition, TRIM56 was proposed to restrict influenza A and B viruses through the C-terminal sequence, independently of the RING, B-box, or coiled-coil domain (39), implying that TRIM56 can restrict evolutionarily divergent viruses through either E3 ligase-dependent or E3 ligase-independent mechanisms.

TRIM69 has been reported to restrict DENV2 through a direct interaction with NS3 coupled with targeting toward proteasomal degradation by ubiquitination at lysine 104 involving its RING domain (40). In contrast, Rihn et al. did not detect TRIM69-mediated restriction of DENV2 NGC in Vero cells and did not observe a reduction in NS3 co-transfected with TRIM69 in HEK293T cells (41). Thus, there is currently very limited knowledge, and no consensus, on whether TRIM69 inhibits DENV2.

Thus, studies published so far suggest that individual TRIM proteins can modulate DENV and other flaviviruses through various mechanisms. However, conflicting results have been reported, and the molecular mechanisms behind TRIM-mediated restriction remain elusive. Also, whether DENV proteins can reciprocally modulate these TRIMs has received little attention. Here, we characterized the effects of TRIM5α, TRIM56, and TRIM69 against DENV2 in a side-by-side manner, using overexpression as well as CRISPR-Cas9-mediated knockout in multiple cell lines in order to generate gain- and loss-of-function data. We also analyzed TRIM-NS2B3 modulation effects in virus-free experiments. Our results reveal novel DENV2-modulating activities for TRIM5α and TRIM69 and suggest TRIM proteins targeting by DENV2 NS2B3.

## RESULTS

### TRIM5α and TRIM56, but not TRIM69, restrict DENV2 infection

TRIM-mediated antiviral activity is virus-, host- and cell context-specific (35, 42). Focusing on the IFN-I-induced cytoplasmic TRIM proteins TRIM5α, TRIM56 and TRIM69, we compared their activity on DENV2 infection in multiple cell types. For this, we used retroviral or lentiviral transduction to stably overexpress each of them in Vero E6, HEK293T, and HeLa cells. Expression of HA-or FLAG-tagged TRIM proteins was confirmed by immunoblotting (Fig. S1), and the cells were challenged with the DENV2 strain 16681s modified to express GFP (43). In this DENV construct, GFP is inserted between the 5’ cyclization sequence and the viral capsid (C) protein gene. GFP is thus expressed concomitantly to the other viral proteins, whereas a self-cleaving 2A peptide sequence placed between GFP and C prevents its expression as a fusion protein. Infection was quantified by flow cytometry based on the percentage of GFP-positive cells.

In Vero E6 cells, representative flow cytometry profiles demonstrated a significantly lower abundance of GFP-positive cells in TRIM5α and TRIM56 expressing cells compared to untransduced control cells (∼5-fold and ∼2-fold, respectively). TRIM69 overexpression, however, resulted in a ∼2.5-fold increase in infected cells (Fig. 1A and B). Upon infection with serial two-fold dilutions of DENV2-GFP, we again observed reduced percentages of GFP-positive cells in cells transduced with TRIM5α or TRIM56 compared to untransduced controls at every dilution examined (Fig. S2A and B). This restriction pattern was recapitulated in HEK293T (Fig. 1C) and HeLa cells (Fig. 1D). In these two cell lines, TRIM5α and TRIM56 transduction consistently reduced DENV2 infection. Co-transduction of TRIM5α and TRIM56 in HEK293T cells showed a higher level of restriction compared to cells transduced with either TRIM5α or TRIM56 (Fig. 1C). In contrast, TRIM69 overexpression led to a modest but significant increase in the proportion of infected cells compared with untransduced controls in both cell lines. In addition to percentages of infected cells, we analyzed the intensity of GFP expression as a marker for viral protein expression levels in infected (GFP-positive) cells (Fig. S3). We found that TRIM5α- and TRIM56-mediated restriction was accompanied by a significant decrease in GFP mean fluorescence intensity in HeLa cells but not in Vero E6 or HEK293T cells. It is possible, however, that GFP expression is at saturating levels in the two latter cell lines, which would explain why TRIM5α and TRIM56 do not significantly downmodulate it. Alternatively, TRIM-mediated restriction in these two cell lines may lead to a total collapse of replication in some cells whereas having little impact in others.

**FIG 1.**
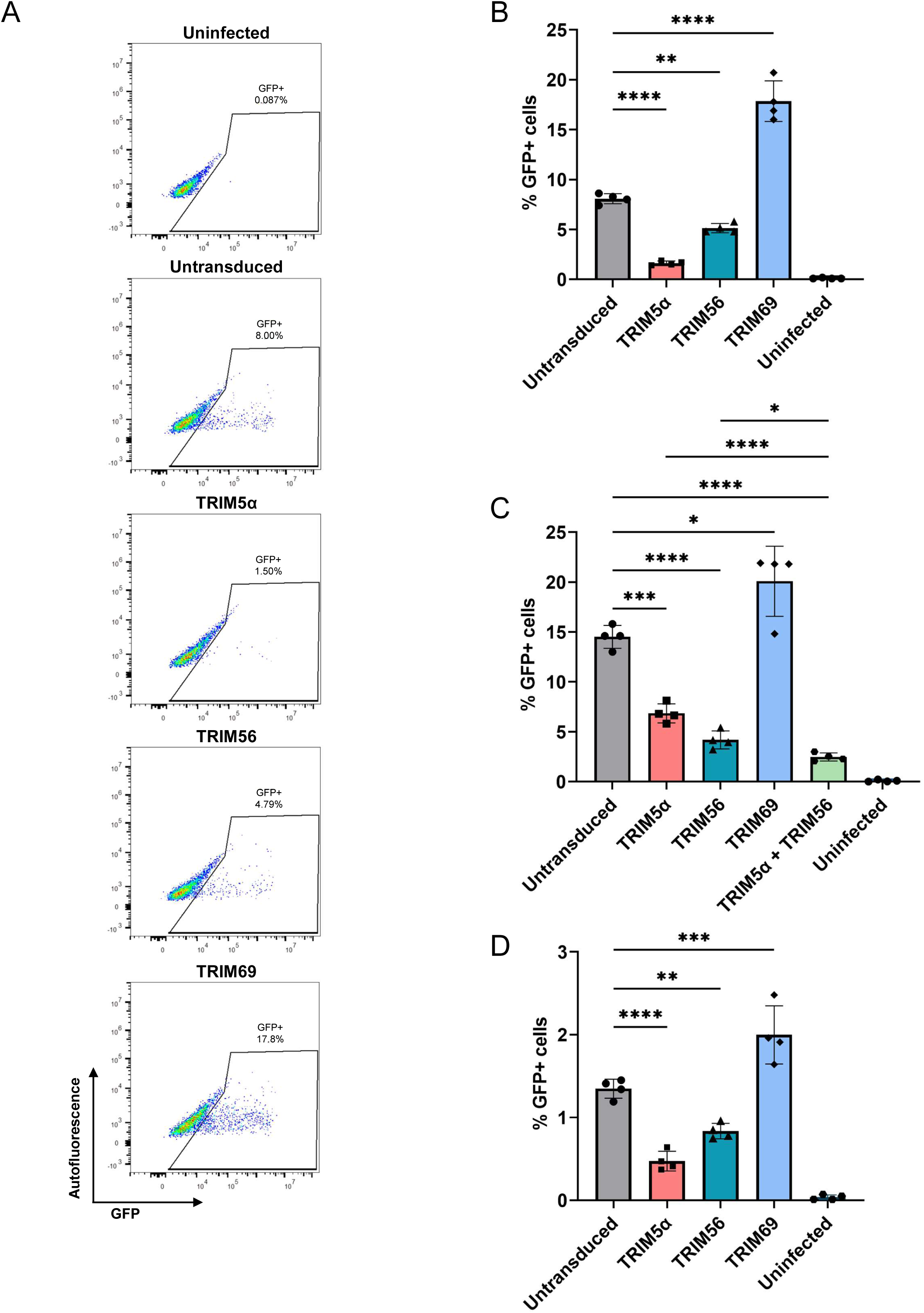
TRIM5α and TRIM56 but not TRIM69 restrict DENV2. (A) Representative pseudo-color dot plots of Vero E6 cells transduced with TRIM5α, TRIM56, or TRIM69, or untransduced control cells infected with DENV2-GFP at a multiplicity of infection (MOI) of 0.05. Cellular autofluorescence (y-axis) is plotted against GFP intensity (x-axis), with the percentages of GFP-positive cells indicated on each plot. (B) Percentage of GFP-positive Vero E6 cells following stable transduction of TRIM proteins and infection with DENV2-GFP as in (A). (C-D) Percentage of GFP-positive HEK293T (C) and HeLa (D) cells stably expressing TRIM5α, TRIM56, TRIM69, or a combination of TRIM5α and TRIM56, or untransduced control cells infected with DENV2-GFP as described in Methods. Data for (B-D) are representative of multiple independent experiments and are from triplicate infections ± SD. ns = non-significant, *P < 0.05, **P < 0.01, ***P < 0.001, ****P < 0.0001, determined by one-way ANOVA with Dunnett’s post-test.

Using GFP as an infection marker in short-term infection experiments may mask restriction mechanisms targeting late replication stages. In order to investigate the effects of TRIM proteins overexpression on infectious virus production, culture supernatants of untransduced or TRIM-transduced Vero E6 cells infected with wild type (WT) DENV2 were titrated for infectious virus yield by plaque assay. Results revealed that supernatants from TRIM5α- and TRIM56-expressing cells yielded less plaques compared to untransduced control cells, in contrast to TRIM69-expressing cells which produced more plaques (Fig. 2A). Quantification confirmed a significant decrease in infectious virus titers in TRIM5α- and TRIM56-expressing cells (∼7-fold and ∼4.3-fold, respectively), contrasting with an increase in TRIM69-transduced cells. Together, these data show that TRIM5α and TRIM56 but not TRIM69 restrict DENV2 infection across three cell lines of primate origin, extending the antiviral spectrum of TRIM5α to DENV2.

**FIG 2.**
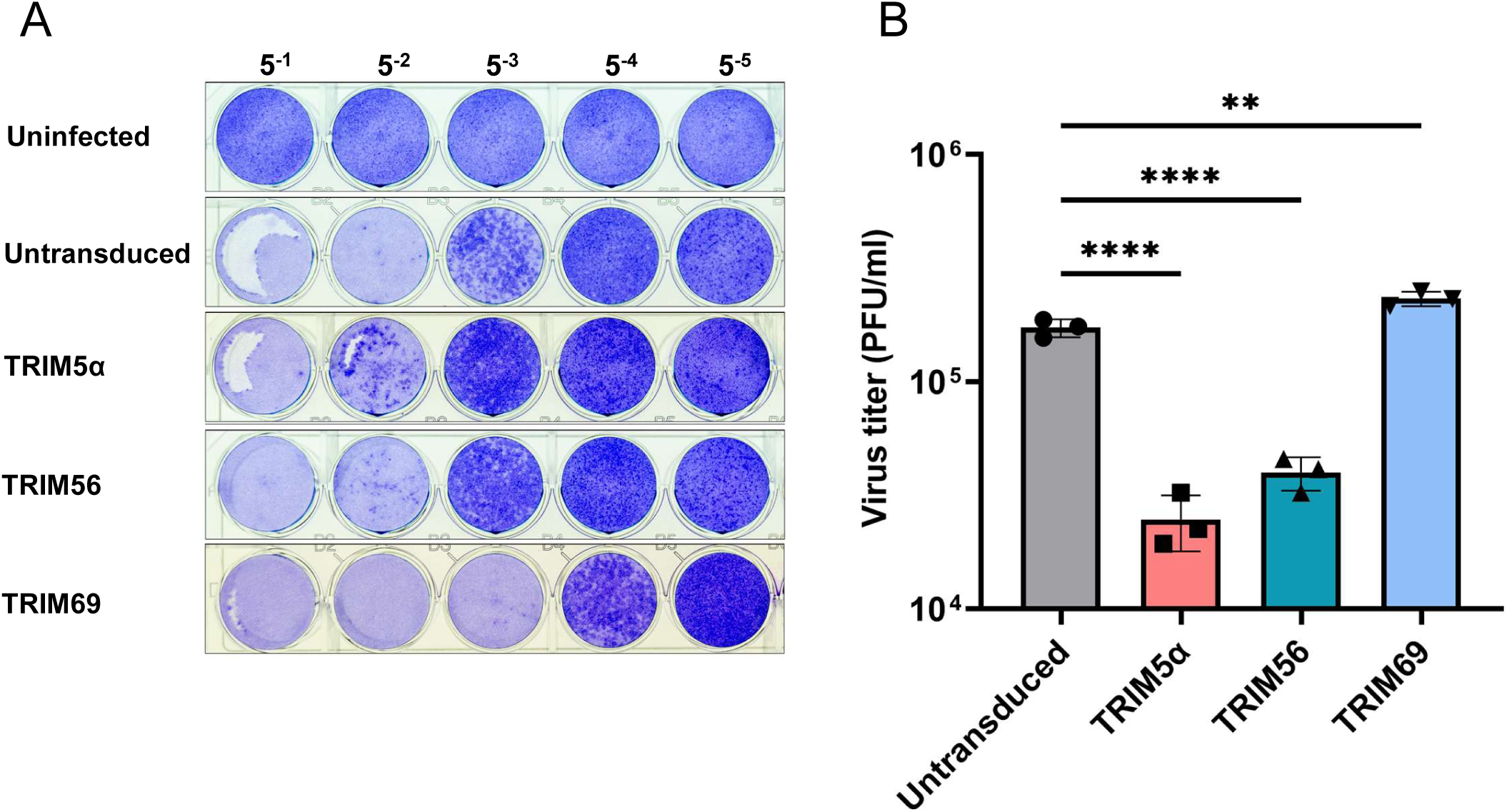
TRIM5α and TRIM56 but not TRIM69 decrease infectious DENV2 production. (A) Representative plaque assay of supernatants collected from untransduced Vero E6 cells, or from cells stably transduced with the indicated TRIMs, and infected with WT DENV2. The plaque assay was performed using serial five-fold dilutions. Supernatant from uninfected cells was included as a negative control. (B) Infectious DENV2 titers expressed as PFU/mL following plaque quantification. Data are from triplicate infections ± SD. ns = non-significant, **P < 0.01, ****P < 0.0001, determined by one-way ANOVA with Dunnett’s post-test.

### TRIM5α and TRIM56 reduce DENV2 proteins and RNA levels

To determine whether the reduction in infection yields resulting from TRIM5α or TRIM56 overexpression was accompanied with a decrease in viral protein and RNA levels, we infected HEK293T cells overexpressing each TRIM with DENV2-GFP for 24 h and analyzed them by immunoblotting. We observed lower levels of the viral NS3 and NS5 proteins in cells transduced with TRIM5α or TRIM56, or co-transduced with both, relative to untransduced cells (Fig. 3A). We next asked whether restriction was also reflected at the level of viral RNA. TRIM-transduced cells were tested for infection using DENV2-GFP, and we observed again that both TRIM5α and TRIM56 reduced HEK293T cell permissiveness to DENV2 infection (Fig. 3B). In parallel, DENV2 RNA was prepared from infected cells and quantified at 24 and 48 hours post-infection (hpi) by qRT-PCR as described in Methods (Fig. 3C-D). Untransduced cells treated with the viral polymerase inhibitor NITD served as a control for viral RNA replication inhibition. At 24 hpi, similar DENV2 RNA levels were observed in untransduced cells and TRIM5α-expressing cells. Although viral RNA levels were lower in TRIM56-expressing cells, this effect did not quite reach statistical significance (P = 0.077) (Fig. 3C). At 48 hpi, both TRIM5α and TRIM56 significantly reduced DENV2 RNA levels relative to untransduced cells, with a greater decrease observed for TRIM56 than for TRIM5α (∼13-fold and ∼5-fold, respectively) (Fig. 3D). Restriction of DENV2 by TRIM5α and TRIM56 is therefore accompanied with a reduction in viral protein and genomic RNA accumulation.

**FIG 3.**
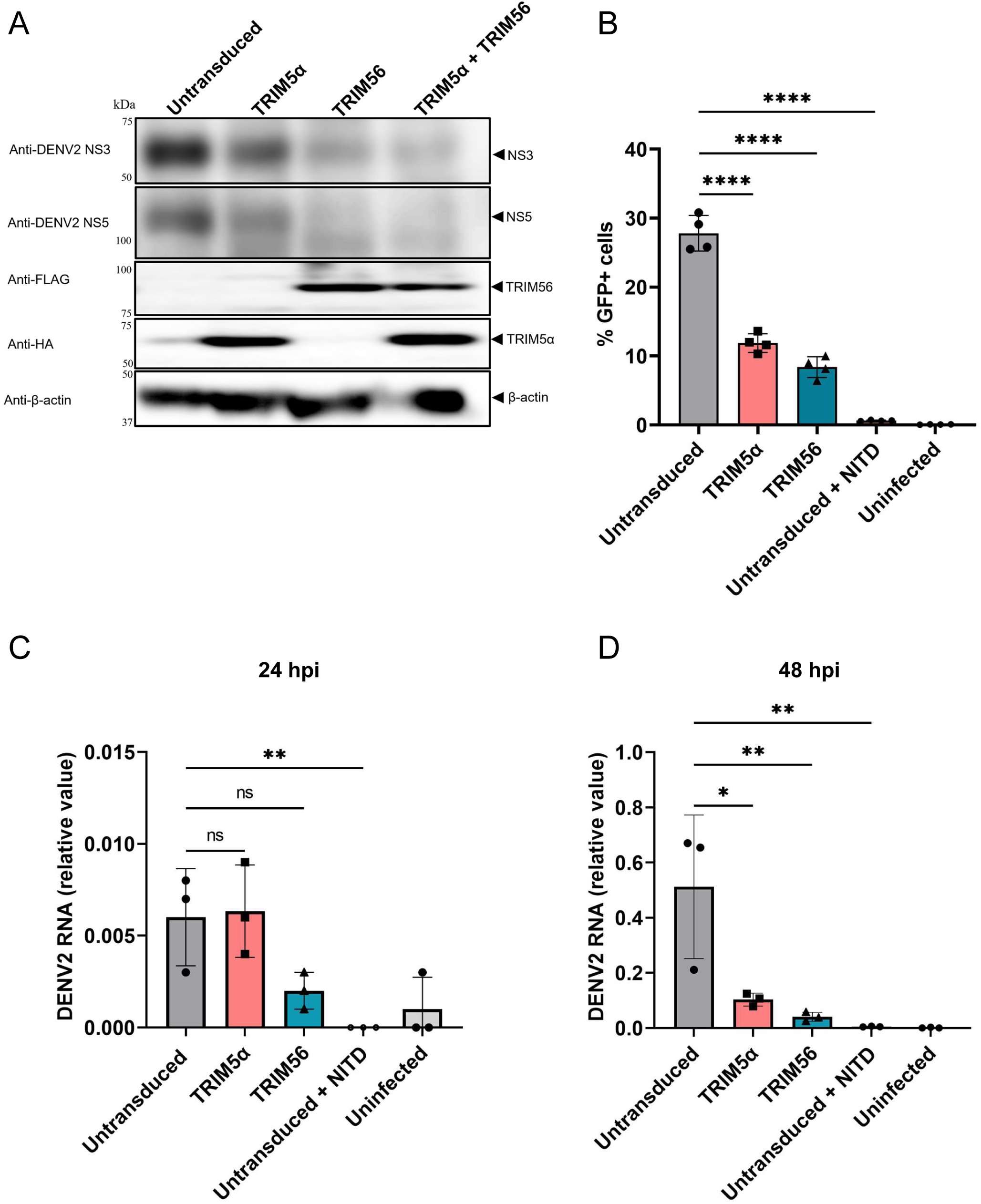
TRIM5α and TRIM56 reduce DENV2 protein levels and viral RNA in HEK293T cells. (A) WB analysis of HEK293T cells stably expressing TRIM5α, TRIM56, or untransduced control, and infected with DENV2-GFP as described in Materials and Methods. Lysates were prepared 24 hpi, processed for WB and probed with antibodies against DENV2 NS3, NS5, FLAG (TRIM56), HA (TRIM5α) and β-actin. Molecular weight markers and the expected positions of the proteins analyzed are indicated. (B-D) Quantification of viral RNA in infected HEK293T cells. (B) Percentage of GFP-positive cells in HEK293T cells transduced with either TRIM5α or TRIM56 or left untransduced, two days after infection with DENV2-GFP in triplicate. Untransduced cells infected in presence of NITD and uninfected cells were also included. (C and D) Relative DENV2 RNA in cells treated as in (B), as determined by qRT-PCR at 24 hpi (C) or 48 hpi (D) using the ΔΔCt method with GAPDH for normalization. ns = non-significant, *P < 0.05, **P < 0.01, ****P < 0.0001, determined by one-way ANOVA with Dunnett’s post-test.

### Endogenous TRIM5 and TRIM56 restrict DENV2 infection, whereas TRIM69 promotes infection

To complement the overexpression approach and to investigate whether the endogenous TRIM5α, TRIM56, and TRIM69 proteins also influence DENV2 infection, we knocked out their corresponding genes by CRISPR/Cas9 in HeLa and Huh7 cells. We targeted TRIM5 using a previously described lentiviral CRISPR construct (“gRNA2”) (44). For TRIM56 and TRIM69, we generated and tested two independent constructs, each expressing a different guide RNA (gRNA). Editing efficiency was validated by Sanger sequencing and TIDE analysis (Fig. S4 A-G). Efficient knockout by indel mutations was detected for TRIM5, as well as for the various gRNAs targeting TRIM56 and TRIM69.

In HeLa cells, knockout of either TRIM5 or TRIM56 resulted in a ∼2-to ∼2.5-fold increase in the percentage of DENV2-GFP positive cells compared with control cells (Fig. 4A), supporting an antiviral role for these two proteins. In contrast, TRIM69 knockout decreased DENV2 infection, suggesting that endogenous TRIM69 acts as a positive factor rather than a restriction factor. A similar pattern was observed in Huh7 cells analyzed by flow cytometry at two different time points (48 and 72 h) (Fig. 4B-C). In these cells, both TRIM5 and TRIM56 knockout resulted in a significant increase in DENV2 infection at both 48 and 72 hpi (up to ∼5-fold for TRIM5, ∼2-fold for TRIM56; Fig. 4B). Conversely, TRIM69 knockout resulted in a decrease in infection at both time points (∼2-to 3-fold; Fig. 4C). Results from these loss-of-function experiments are therefore consistent with the overexpression approach (Fig. 1 and 2) and identify TRIM5α and TRIM56 as restriction factors for DENV2, while showing a proviral role for TRIM69.

**FIG 4.**
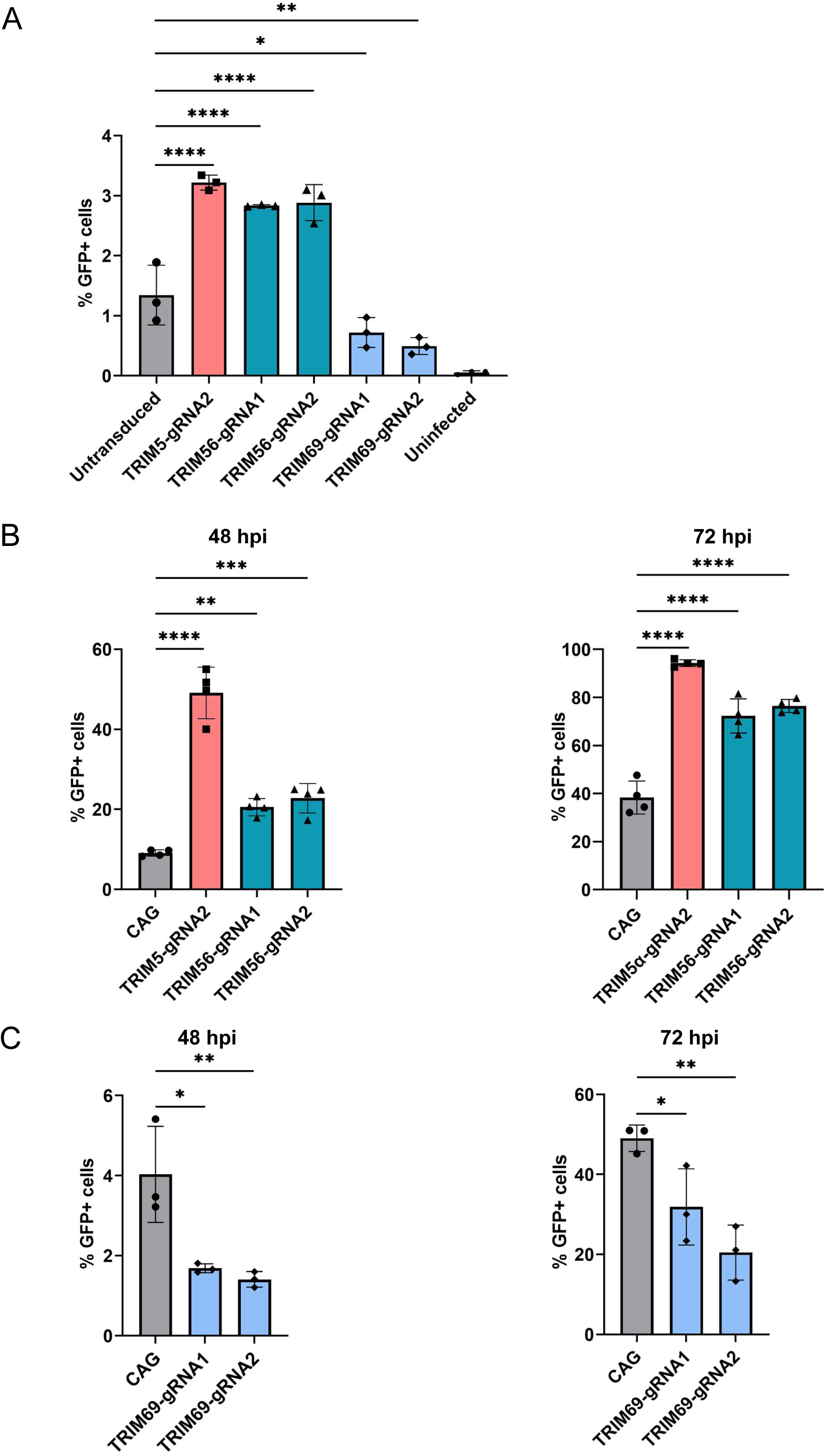
CRISPR knockout of TRIM5 and TRIM56, but not TRIM69, increases susceptibility to DENV2 infection. (A) Percentage of GFP-positive cells determined by flow cytometry in HeLa cells transduced or not with CRISPR-Cas9 constructs targeting TRIM5, TRIM56 or TRIM69, and infected with DENV2-GFP for two days. (B) Percentage of DENV2-GFP-positive cells in control (CAG-targeting CRISPR) Huh7 cells, or in cells expressing TRIM5- or TRIM56-targeting gRNAs, following DENV-GFP infection at 48 hpi (left) and 72 hpi (right). (C) Percentage of DENV2-GFP-positive Huh7 cells transduced with CRISPR constructs expressing a CAG-targeting control gRNA or gRNAs targeting TRIM69 at 48h (left) and 72 h (right) following DENV-GFP infection. Data are from three (A and C), and four (B) independent experiments ± SD. ns = non-significant, *P < 0.05, **P < 0.01, ***P < 0.001, ****P < 0.0001, determined by one-way ANOVA with Dunnett’s post-test, with comparisons done relative to the untransduced (A), and CAG (B and C) controls.

### Reciprocal downmodulation of TRIM56 and DENV2 NS2B3 independently of NS2B3 protease catalytic activity

We next performed virus-free transfection experiments to investigate whether TRIM proteins might affect essential DENV2 replication enzymes, and reciprocally, whether DENV2 proteins might modulate TRIM proteins. In particular, the flaviviral NS2B3 heterodimer was proposed to be the molecular target of restriction factors TRIM5α and TRIM69 (10). Conversely, NS2B3 is also known to modulate host antiviral proteins through both protease-dependent and protease-independent mechanisms (40, 45–47). Thus, HEK293T cells were co-transfected with the WT NS2B3 (NS2B3-WT) or the catalytically dead mutant NS2B3-S135A together with each TRIM protein (Fig. 5A), and we used densitometry of WB signals to analyze TRIM and NS2B3 protein levels (Fig. 5B-G).

**FIG 5.**
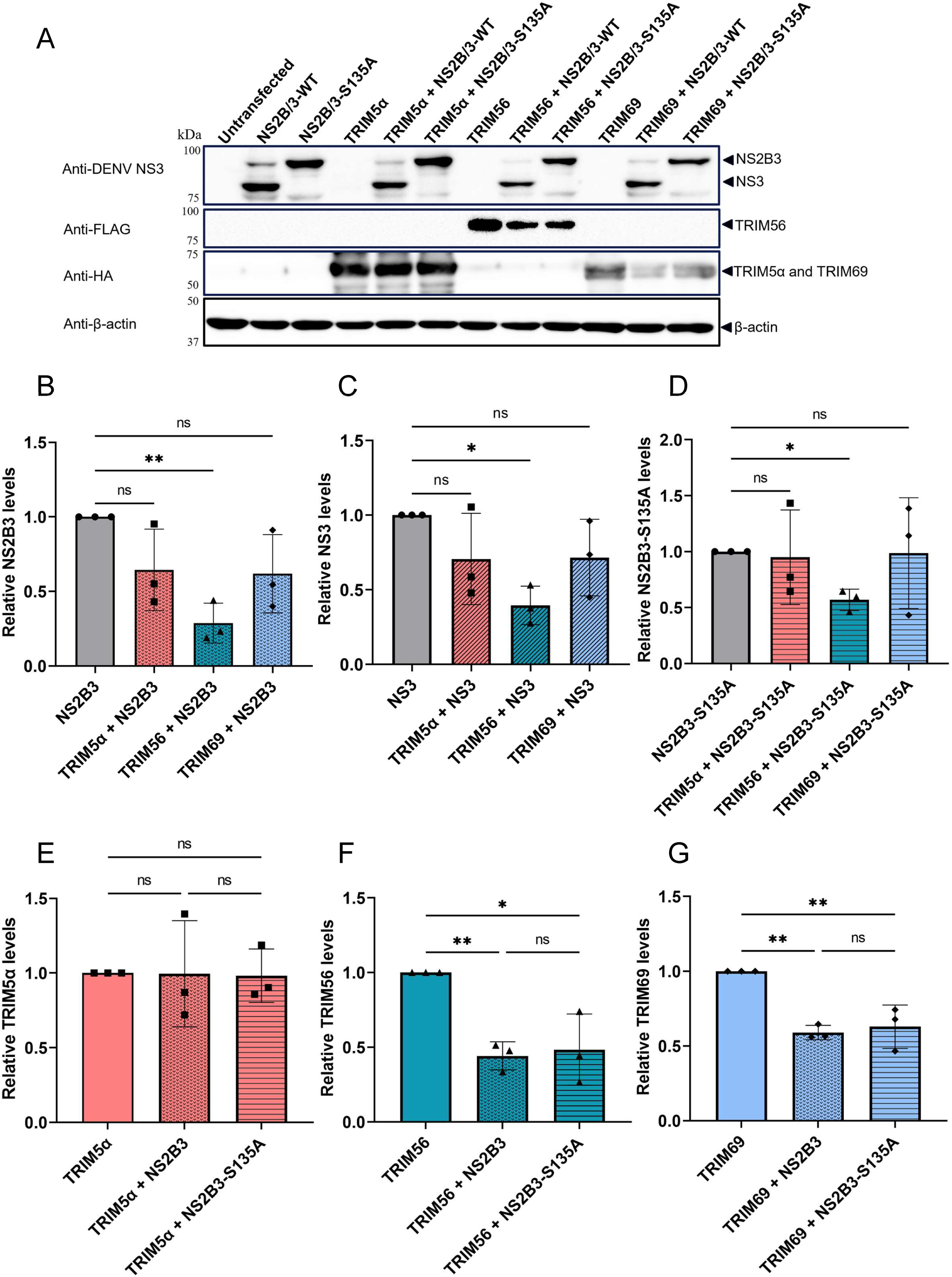
Reciprocal effects between TRIM proteins and DENV2 NS2B3 in virus-free expression assays. (A) WB analysis of HEK293T cells transfected or not with DENV2 NS2B3-WT or its protease-defective NS2B3-S135A mutant, and with TRIM5α, TRIM56 or TRIM69. Detection was done using antibodies against DENV NS3 (detecting both the NS2B3 precursor and cleaved NS3 product), FLAG (TRIM56), HA (TRIM5α and TRIM69), and β-actin. (B-D) Densitometric quantification of NS2B3-WT (B), cleaved NS3 (C), and NS2B3-S135A (D) levels relative to β-actin, normalized to the corresponding construct expressed alone. (E-G) Densitometric quantification of TRIM5α (E), TRIM56 (F), and TRIM69 (G) levels relative to β-actin, normalized to each TRIM expressed alone. Data are from three independent experiments ± SD. ns = non-significant, *P < 0.05, **P < 0.01 by one-way ANOVA with Dunnett’s (B-D) and Tukey’s post-test (E-G).

We found that TRIM56 significantly reduced NS2B3-WT levels (Fig. 5B), whereas TRIM5α and TRIM69 had a smaller effect that did not reach statistical significance in this context (Fig. 5B). We were also able to analyze the NS3 cleaving product and found that once again, TRIM56 but not TRIM5α nor TRIM69 significantly decreased its abundance (Fig. 5C). TRIM56 also significantly reduced the catalytically inactive NS2B3-S135A mutant, whereas TRIM5α and TRIM69 had no effect on this protein (Fig. 5D). In summary, TRIM56 was the only TRIM protein able to significantly affect NS2B3 levels in these conditions, and this phenotype was not dependent on NS2B3 catalytic activity.

TRIM5α levels remained unchanged in presence of either WT NS2B3 or the S135A mutant (Fig. 5E). However, TRIM56 and TRIM69 levels were each significantly reduced by WT NS2B3, and a similar decrease was observed with the S135A mutant (Fig. 5F and 5G). Thus, DENV2 NS2B3 selectively reduces TRIM56 and TRIM69 levels, but not TRIM5α levels, and this effect does not require the NS2B3 protease catalytic activity. Altogether, these data point to a reciprocal modulation between TRIM56 and the DENV2 NS2B3 complex.

### TRIM56 and TRIM69 levels are reduced by the NS2B3 complex but not by NS3 alone

We next analyzed whether the mature NS3 protease, independent of its NS2B cofactor, was sufficient to downmodulate TRIM56 and TRIM69. TRIM5α, TRIM56, or TRIM69 were therefore co-transfected together with either NS2B3 or NS3, and TRIM protein levels were analyzed by immunoblotting (Fig. 6A) followed by densitometry (Fig. 6B-D). As expected, TRIM5α protein levels were not significantly altered by either NS2B3 or NS3 alone (Fig. 6B). As previously observed, NS2B3 significantly reduced TRIM56 levels (Fig. 6C) as well as TRIM69 levels (Fig. 6D). Although reduced TRIM56 levels were also observed in the presence of NS3, this effect was not statistically significant compared with TRIM56 expressed alone (Fig. 6C). Similar results were obtained for TRIM69 (Fig. 6D).

**FIG 6.**
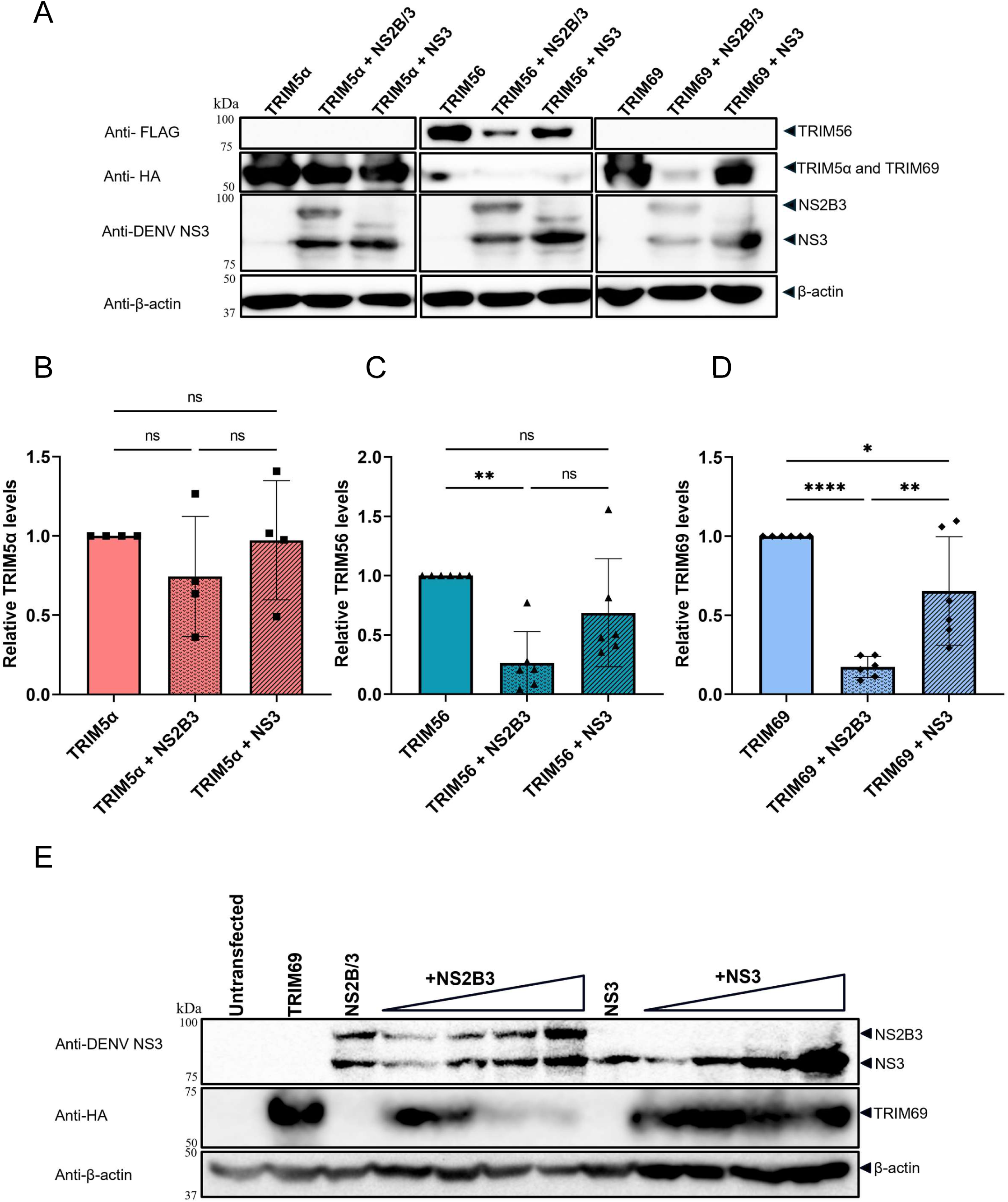
DENV2 NS2B3 but not NS3 reduce TRIM56 and TRIM69 levels. (A) WB analysis of HEK293T cells co-transfected with NS2B3 or NS3 and with TRIM5α, TRIM56, or TRIM69, or transfected with each construct alone, probed with antibodies against FLAG (TRIM56), HA (TRIM5α and TRIM69), DENV NS3 (detecting both the NS2B3 precursor and NS3 mature product), and β-actin. (B-D) Densitometric quantification of TRIM5α (B), TRIM56 (C), and TRIM69 (D) levels relative to β-actin, normalized to each TRIM expressed alone. (E) WB analysis of HEK293T cells co-transfected with a fixed amount of TRIM69 and increasing amounts of NS2B3 or NS3, probed with antibodies against DENV NS3, HA (TRIM69), and β-actin. WBs were analyzed using the indicated antibodies. Data are from four (B) or six (C-D) independent experiments ± SD. ns = non-significant, *P < 0.05, **P < 0.01, ****P < 0.0001 by one-way ANOVA with Tukey’s post-test.

To further investigate the different effects of NS2B3 and NS3 on TRIM69, we co-transfected increasing amounts of either NS2B3 or NS3 with TRIM69. A dose-dependent reduction in TRIM69 levels was observed with increasing amounts of NS2B3, whereas NS3 had no effect (Fig. 6E). Altogether, the results from Fig. 6 suggest that the whole NS2B3 complex, rather its NS3 subunit, is required to mediate TRIM proteins downmodulation, pointing to the importance of correct NS2B3 localization and folding for this function.

### The role of TRIM56 RING domain in DENV2 restriction and TRIM56-NS2B3 reciprocal modulation

TRIM56 antiviral activity against flaviviruses, including DENV2 and ZIKV, has been reported to depend on its RING domain (37, 38). In order to investigate the role of the RING domain in DENV2 restriction and in TRIM56-NS2B3 interactions, we used the RING domain mutant CC21/24AA (37) abrogating ubiquitin ligase activity (TRIM56-RINGmut). We first compared WT TRIM56 and TRIM56-RINGmut for their ability to restrict DENV2-GFP infection (Fig. 7A-B). In HEK293T cells, transduction of WT TRIM56 significantly lowered DENV2-GFP infection compared to untransduced cells (3.3-fold), as expected. TRIM56-RINGmut showed only partial restriction activity (1.4-fold), which was significantly weaker than that of the WT TRIM56 but not abrogated (Fig. 7A). In Vero E6 cells, both WT and RING-mutated TRIM56 efficiently restricted DENV2 infection (Fig. 7B), with the RING mutant showing a smaller level of inhibition compared to the WT (4.1-fold and 2.5-fold, respectively) but without reaching statistical significance. These results show that the RING domain contributes to TRIM56-mediated restriction, but is not essential, and suggest a cell context-specific effect.

**FIG 7.**
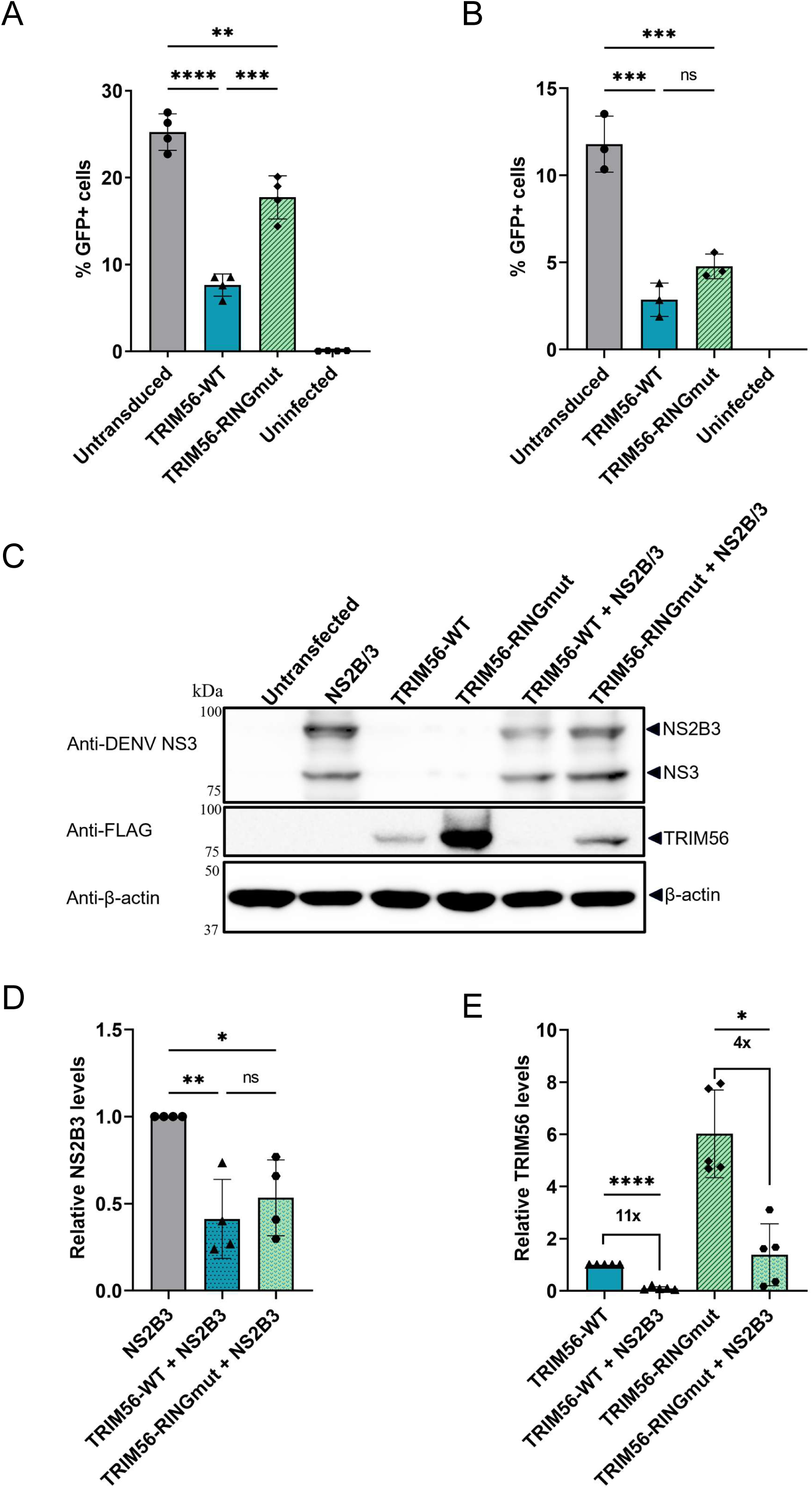
TRIM56-NS2B3 reciprocal downmodulation does not require TRIM56 E3 ubiquitin ligase activity. (A and B) Percentage of GFP-positive HEK293T cells (A) and Vero E6 cells (B) stably transduced with WT TRIM56 or its RING domain mutant CC21/24AA (RINGmut), or left untransduced, and infected with DENV2-GFP for two days. Shown are means from infections done in quadruplicate (A) or triplicate (B), with SD. (C) WB analysis of HEK293T cells co-transfected with NS2B3 and with WT TRIM56 or TRIM56-RINGmut, or transfected with each construct alone, probed with antibodies against DENV NS3, FLAG (TRIM56), and β-actin. (D and E) Densitometric quantification of NS2B3 levels (D) and TRIM56 levels (E) relative to β-actin and normalized to NS2B3 or TRIM56-WT expressed alone, respectively. Data are from four (D) or five (E) independent experiments ± SD. ns = non-significant, *P < 0.05, **P < 0.01, ***P < 0.001, ****P < 0.0001 by one-way ANOVA with Tukey’s post-test.

We next investigated whether the RING mutation affected the reciprocal modulation of TRIM56 and NS2B3 in virus-free assays (Fig. 7C-E). WT TRIM56 or its RING mutant were transfected alone or with NS2B3 in HEK293T cells followed by immunoblotting (Fig. 7C). Densitometric quantifications showed a significant reduction in NS2B3 levels in the presence of either WT or mutant TRIM56, with no significant difference between WT and RING-mutated TRIM56, demonstrating that the RING domain is not required for this specific phenotype (Fig. 7D). RING-mutated TRIM56 accumulated at significantly higher levels than WT TRIM56 when transfected alone (∼6-fold), suggesting RING-mediated self-ubiquitination (48). Co-transfection with NS2B3 significantly lowered the levels of both WT and RING mutant TRIM56, reaching an approximately 11-fold decrease for the former and 4-fold for the latter (Fig. 7C and E). Together, these results suggest that the TRIM56 RING mutant retains the ability to reduce DENV2 NS2B3 levels, despite a diminished restriction activity in HEK293T cells. Likewise, NS2B3 downmodulates both WT and RING-mutant TRIM56, though it may have a weaker effect on the RING mutant. Hence, the TRIM56 RING E3 ubiquitin ligase activity is not essential for the observed reciprocal downmodulation between TRIM56 and NS2B3.

### NS2B3-mediated downmodulation of TRIM56 and TRIM69 and the reciprocal down-modulation of NS2B3 by TRIM56 are not rescued by proteasome or lysosome inhibition

Both proteasomal and lysosomal pathways have previously been proposed to be involved in TRIM-mediated antiviral activity. For instance, TRIM5α was reported to bind the NS2B3 protease complex of Langat virus (LGTV), promoting its K48-linked ubiquitination and proteasomal degradation (35), and has also been linked to autophagy and lysosomal pathways upon HIV-1 restriction (49, 50). On the other hand, DENV NS2B3 can induce lysosome-dependent degradation of the host factor Nrf2 (47). Therefore, we investigated whether inhibition of the proteasomal and lysosomal pathways could counteract TRIM-mediated restriction or the TRIM-NS2B3 reciprocal downmodulation effects. HEK293T cells expressing TRIM56 or TRIM69 alone or co-expressed with NS2B3 were treated with the proteasome inhibitor MG132 or the lysosomal inhibitor bafilomycin A1 (BafA1) for 6 h prior to lysis, followed by immunoblot analysis (Fig. 8A) and densitometry analyses (Fig. 8B-D). Neither inhibitor affected NS2B3 levels when NS2B3 was transfected alone (Fig. 8B). Consistent with our earlier results, TRIM56 co-transfection lowered NS2B3 levels; however, neither inhibitor significantly restored NS2B3 levels in the presence of TRIM56 (Fig. 8B). MG132 significantly increased TRIM56 levels in the absence of NS2B3, whereas BafA1 had no significant effect (Fig. 8C), suggesting that TRIM56 is subjected to basal proteasomal degradation under these conditions. As observed previously, TRIM56 levels were lowered when co-transfected with NS2B3, but neither MG132 nor BafA1 significantly restored TRIM56 levels (Fig. 8C). Similarly, neither inhibitor significantly modulated TRIM69 levels when transfected alone, and neither treatment reversed the significant decrease in TRIM69 induced by NS2B3 co-transfection (Fig. 8D).

**FIG 8.**
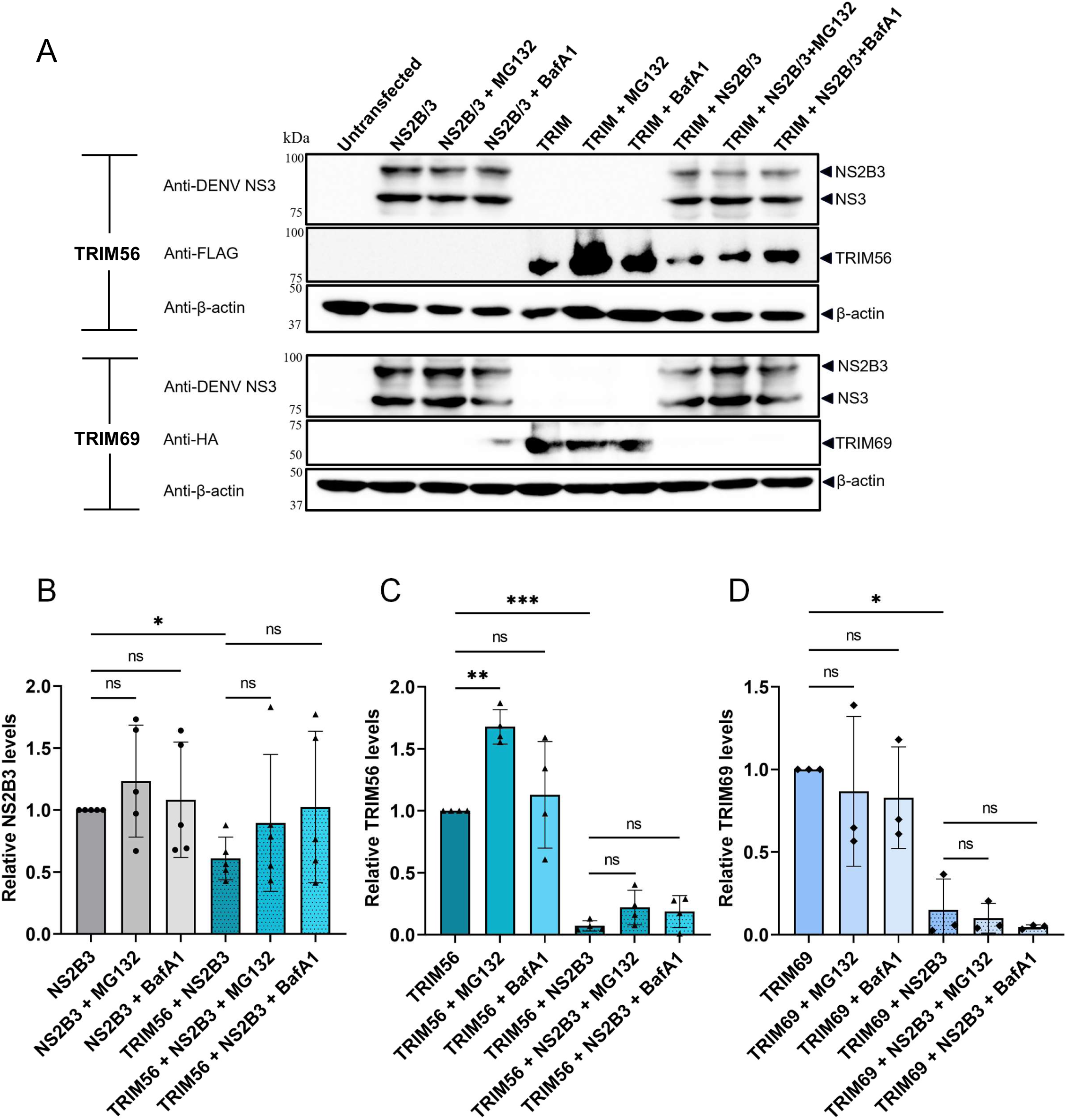
NS2B3-mediated downmodulation of TRIM56 and TRIM69, and TRIM56-mediated downmodulation of NS2B3, are not rescued by proteasome or lysosome inhibition. (A) WB analysis of HEK293T cells co-transfected with NS2B3 and with TRIM56 or TRIM69, or transfected with each construct alone, treated with the proteasome inhibitor MG132 (10 µM) or the lysosomal inhibitor bafilomycin A1 (BafA1) (1 µM) for 6 h before lysis at 48 h post-transfection, and probed with antibodies against DENV NS3, FLAG (TRIM56) or HA (TRIM69), and β-actin. (B-D) Densitometric quantification of NS2B3 (B), TRIM56 (C), and TRIM69 (D) levels, relative to β-actin and normalized to each construct expressed alone. Data are from four (B and C) and three (D) independent experiments ± SD. ns = non-significant, *P < 0.05, **P < 0.01, ***P < 0.001 by one-way ANOVA with Tukey’s post-test.

We also investigated whether MG132 would modulate DENV2-GFP infection of TRIM5α-or TRIM56-transduced cells (Fig. S5). This analysis was complicated by the cytotoxic effects of the drug, leading us to use lower concentrations than were used in the virus-free assays. HEK293T cells transduced or not with TRIM5α or TRIM56 were infected with DENV2-GFP at multiple dilutions in the presence or absence of MG132 at two different concentrations (Fig. S5). In the conditions tested, MG132 did not detectably modulate DENV2-GFP infection in either TRIM5α-or TRIM56-expressing cells (Fig. S5). Together, our findings suggest that the TRIM-NS2B3 reciprocal downmodulation effects are independent of either the proteasomal or lysosomal pathways. Consistent with this, proteasome inhibition did not reverse the antiviral phenotype of TRIM5α or TRIM56 during DENV infection.

## DISCUSSION

TRIM proteins have emerged as key modulators of viral replication, functioning either as intrinsic cellular restriction factors or as regulators of antiviral signaling pathways (10, 22). Their activity can vary depending on the virus and the cellular background, as shown by cell type-dependent antiviral phenotypes reported for individual TRIM proteins (41, 42, 51). Here, we investigated three cytoplasmic TRIM proteins that have been previously proposed to modulate flavivirus infection. TRIM56 was identified as a restriction factor for DENV2 by one research group (37). TRIM5α was reported not to inhibit DENV2 in another study (35), whereas previous studies on TRIM69 have led to inconsistent conclusions (40, 41). In our study, TRIM5α and TRIM56 restricted DENV2 strain 16681, whereas TRIM69 facilitated infection. These observations were made using both stable overexpression and endogenous loss-of-function approaches. Restriction by TRIM5α and TRIM56 was accompanied with reduced accumulation of NS3 and NS5 and, at 48 hpi, lower levels of viral RNA were observed. Virus-free co-expression experiments further demonstrated reciprocal downmodulation between TRIM56 and NS2B3 proteins, and NS2B3 also reduced TRIM69 levels.

Our data establish human TRIM5α as a restriction factor for DENV2 strain 16681. TRIM5α expression reduced infection in multiple cell lines and decreased infectious-virus production in Vero E6 cells, whereas disruption of endogenous TRIM5 increased infection in both HeLa and Huh7 cells (Figs. 1, 2, and 4). These results are in apparent conflict with the study from Chiramel and colleagues (35), however, the discrepancy might be explained by several key differences in methodology: (i) their study used a different DENV2 strain (NGC); (ii) overexpression experiments were performed with rhesus TRIM5α instead of human as in the present study; (iii) TRIM5 knockout experiments were done in a different cell line (HAP1) (35). Laboratory propagation of DENV strains can introduce additional sequence variation that may potentially modulate sensitivity to restriction factors (52). Recently, it was reported that a single amino-acid substitution in Powassan virus NS3 was sufficient to increase resistance to otherwise restrictive primate TRIM5α proteins, providing direct evidence that limited viral sequence variation can alter flavivirus susceptibility to TRIM5α (36). Conclusively investigating the role of DENV strain specificity in TRIM5α-mediated restriction will require the direct comparison of DENV2 NGC and 16681 in infection assays. Cellular context may also contribute, since TRIM-associated antiviral phenotypes can vary among cell types (42). In our study, however, the phenotype was reproduced in several cell lines and by both gain- and loss-of-function approaches, arguing against an effect confined to a single cellular background.

The mechanism by which TRIM5α restricts DENV2 remains unresolved. TRIM5α can couple direct recognition of viral structures to activation of innate immune signaling (33). Our experiments do not distinguish whether DENV2 restriction results from the direct inhibition of viral replication, enhanced antiviral signaling, or contributions from both. At 24 hpi, TRIM5α expression was associated with reduced NS3 and NS5 levels without a detectable decrease in viral RNA, whereas viral RNA was significantly reduced at 48 hpi (Fig. 3). This temporal pattern does not identify the primary step affected and could reflect an antiviral effect that becomes amplified as infection proceeds, consistent with the activation of innate pathways. IFN-I production and innate immune signaling were not analyzed, and therefore this possibility remains to be tested. However, IFN-I is not being produced in Vero E6 cells due to a genomic deletion (53), supporting a direct anti-DENV2 role for TRIM5α. The broad antiviral spectrum of TRIM5α may reflect its capacity to recognize different viral determinants. In addition to the retroviral capsid lattice and the NS2B3 complex of susceptible flaviviruses, human TRIM5α has been shown to recognize the orthopoxvirus capsid protein L3 and, more recently, the HSV-1 capsid protein VP19C (34, 35, 54). Determining whether a DENV2 structural protein, such as the nucleocapsid protein, is directly targeted by human TRIM5α will require direct protein-protein interaction evidence, or the identification of DENV2 variants that modulate its sensitivity to TRIM5α. Our virus-free experiments do not hint at DENV2 NS2B3 as being targeted to degradation by TRIM5α. TRIM5α did not significantly reduce the abundance of WT NS2B3, processed NS3, or catalytically inactive NS2B3-S135A (Fig. 5). This differs from the reduction of NS2B3 reported for TRIM5α-sensitive tick-borne flaviviruses (35), but does not exclude the possibility of direct TRIM5α-NS2B3 interactions.

Our findings confirm TRIM56 as a restriction factor for DENV2, consistent with earlier observations of TRIM56-mediated inhibition of DENV2 and YFV RNA replication as reported by Liu et al. (37). In our study, TRIM56 overexpression restricted DENV2 in three different cell lines, whereas knockout of endogenous TRIM56 increased infection in HeLa and Huh7 cells (Figs. 1 and 4). TRIM56 expression was also associated with reduced NS3 and NS5 protein levels and lower viral RNA levels (Fig. 3). These observations are consistent with inhibition of viral RNA replication, although our experiments do not establish the primary stage of restriction. Our results differ, however, with respect to the requirement for the RING domain. Liu et al. reported the loss of DENV2 restriction with the CC21/24AA mutant and concluded that TRIM56 E3 ligase activity was required (37). In our experiments, the same mutation retained measurable antiviral activity in HEK293T cells and Vero E6 cells despite the loss of RING-dependent E3 ligase activity (Fig. 7 A-B). Furthermore, in virus-free experiments, RING-mutated TRIM56 downmodulated NS2B3 as efficiently its WT counterpart, and was itself downmodulated by NS2B3, albeit this effect was smaller compared to WT TRIM56 (Fig. 7 C-E). However, the substantially higher steady-state abundance of TRIM56-RINGmut (Fig. 7C and E) prevents a quantitative assessment of the RING domain’s contributions. Differences in cellular environment or protein expression levels might contribute to the discrepancy with the prior study. Interestingly, RING-independent restriction has been reported against influenza A and B viruses (39), illustrating the possibility of RING domain-independent antiviral activity. We also found that proteasome inhibition did not reverse TRIM56-mediated DENV2 restriction, supporting the RING-independent restriction hypothesis (Fig. S5). However, this finding should be interpreted cautiously because MG132 cytotoxicity, as well as its inhibitory effects on DENV2 infection, prevented us from using it at high concentrations in infection assays.

Our results suggest that the observed TRIM56-dependent NS2B3 protein levels downmodulation is not dependent on proteasomal degradation, nor does it involve the NS2B3 protease activity. Thus, this phenotype might stem from altered protein synthesis, or from decreased protein stability through another pathway. It could also be related to changes in intracellular localization. Determining whether TRIM56 alters NS2B3 RNA abundance, protein half-life, or subcellular localization will be necessary to distinguish among these possibilities.

In contrast to TRIM5α and TRIM56, TRIM69 consistently increased DENV2 infection in our experimental systems. Indeed, TRIM69 overexpression increased infection in all three cell lines tested, whereas disruption of endogenous TRIM69 reduced infection in both HeLa and Huh7 cells (Fig. 1 and 4). This stands in contrast to a previous study reporting that TRIM69 restricts DENV2 by promoting ubiquitination and proteasomal degradation of NS3 (40). However, a recent study disputed those conclusions, showing the absence of TRIM69-mediated DENV2 restriction in Vero E6 cells and no reduced NS3 levels upon co-transfection with TRIM69 in HEK293T cells (41). Whether the absence of a consensus on the role of TRIM69 in DENV infection is explained by differences in the experimental systems used, such as virus strain or protein expression levels, remains to be determined. The positive effect of TRIM69 on DENV2 infection may be explained by interference with other TRIMs. Indeed, heteromerization among virus restriction factors of the TRIM family is experimentally established. For example, TRIM16 can form heteromeric complexes with TRIM18/MID1, TRIM19/PML, and TRIM24 (55), and a more recent work reported that TRIM51 interacts with TRIM49 through their coiled-coil domains and suppresses TRIM49 ubiquitin ligase activity (21). Such interactions may modulate the localization, stability, or activity of individual TRIM proteins. It is therefore possible that the pro-viral role of TRIM69 towards DENV2 infection may be due to TRIM69 negatively modulating another antiviral TRIM protein(s) through heterodimerization. Another possible explanation for the TRIM69 phenotype obtained here is a direct pro-viral activity, perhaps involving the TRIM69 RING domain-mediated ubiquitination of a cellular or viral target. Along those lines, TRIM7 promotes ZIKV entry through K63-linked ubiquitination of the viral envelope protein (56). Determining the DENV2 replication step enhanced by TRIM69 and whether this phenotype depends on its RING domain will be critical to fully understand the mechanism involved.

DENV2 NS2B3 reduced the levels of TRIM56 and TRIM69, whereas TRIM5α remained unaffected (Fig. 5 and 6). Interestingly, both WT NS2B3 and its catalytically inactive S135A mutant significantly reduced TRIM56 and TRIM69 levels, indicating that the NS2B3 protease activity is not required for this effect. This differs from the protease-dependent cleavage of host factors such as STING and cGAS by DENV NS2B3 (57, 58). Our results do not point to a precise mechanism by which DENV NS2B3 downmodulates TRIM56 and TRIM69. Nevertheless, protease-independent modulation of host proteins has been described previously. Ferrari et al. have reported that both WT and S135A NS2B3 reduced Nrf2 protein levels through lysosomal degradation rather than direct proteolysis (47), while Aguirre et al. showed that NS2B expressed alone, but not NS3, was sufficient to promote autophagy via lysosome-dependent degradation of cGAS (59). In addition, DENV2 NS3 expressed alone has also been reported to reduce intracellular RIPK1 levels independent of its protease activity (60). In our hands, NS3 expressed alone did not substantially reduce TRIM56 or TRIM69 levels (Fig. 6). The requirement for NS2B3 in the observed modulation of TRIM proteins could be related to NS2B binding NS3 and contributing to both its conformation and endoplasmic reticulum membrane anchoring (7, 9). A similar dependence on NS2B3 was observed for FAM134B cleavage (61). NS2B3 may also act indirectly by recruiting host-cell proteins that influence TRIM protein stability or degradation. Along those lines, Shah et al. used affinity-purification mass spectrometry to identify interactions between DENV2 NS2B3 and several host proteins belonging to the CTLH complex, a multiprotein complex shown to possess E3 ubiquitin-ligase activity (62, 63).

Together, these studies along with our results suggest that DENV2 NS2B3 can modulate host protein levels independently of its catalytic activity. In our hands, neither MG132 nor bafilomycin A1 rescued NS2B3-mediated TRIM56 or TRIM69 downmodulation (Fig. 8), leaving the mechanism for this decrease in protein levels unresolved. Nonetheless, the observed NS2B3-dependent TRIM reduction may have functional significance as a viral countermeasure against restriction factors. Further investigations will be required to determine whether NS2B3 acts directly on these proteins or through cellular cofactors and why TRIM5α was insensitive to this effect.

Dengue is spreading geographically and the use of the presently available vaccines is restricted to defined populations and in endemic areas with high dengue transmission (64). At present, there is no specific antiviral treatment for dengue, and clinical management remains largely supportive, highlighting the need to characterize viral and host factors that may reveal new therapeutic targets (65). Our findings expand the role of TRIM proteins against DENV2 and show that closely related cytoplasmic TRIM proteins can have opposing effects on viral replication. Our findings broaden the importance of TRIM-mediated regulation of DENV replication and identify host and viral protein regulation pathways that may be explored as possible therapeutic targets. More broadly, understanding how host restriction factors restrict DENV replication and how the virus modulates these defenses may help to identify host-directed or virus-directed strategies for dengue prevention and treatment.

## MATERIALS AND METHODS

### Cell Lines

Human embryonic kidney (HEK) 293T, Hepatocyte-derived carcinoma Huh7, HeLa and African green monkey kidney isolated Vero E6 cells were maintained in Dulbecco’s Modified Eagle Medium (DMEM; Cytiva Hyclone, Logan, UT, USA) supplemented with 10% heat-inactivated fetal bovine serum (FBS; Hyclone) and 1% penicillin and streptomycin (Hyclone) at 37°C and 5% CO2.

### Plasmids

pLVX-huTRIM5a-HA was obtained from Joseph Sodroski (Harvard Medical School, Boston, MA, USA). pMIH-FLAG-huTRIM5α was previously described (66). pCX4bsr-TRIM56-FLAG-WT and the CC21/24AA (RING mutant) version were from Kui Li (University of Tennessee, Memphis, TN, USA) (48). pMIP constructs encoding C-terminally HA-tagged TRIM69 were constructed by transferring the TRIM69 coding sequences from pCR3.1-TRIM69-mCHERRY (67). For this, TRIM69 was PCR amplified using primers (Table S1) incorporating XhoI and EcoRI restriction sites and a HA tag at the C terminus. Following purification, amplicons were digested with XhoI and EcoRI and ligated to pMIP (68) cut with the same enzymes to generate pMIP-TRIM69-HA. The lentiviral expression vector pLentiCRISPRv2 (pLCv2), a gift from Feng Zhang (Addgene #52961) (69), and its blasticidin- and hygromycin-resistant derivatives, plentiCRISPRv2-blast (Addgene #98293) and plentiCRISPRv2-hygro (Addgene #98291) (70) were used for CRISPR/Cas9-mediated knockout of TRIM genes. The plasmid pFK-DVs-G2A encoding DENV2-GFP vector from DENV serotype 2 (DENV2) strain 16681s (GenBank NC_001474) was gifted by Ralf Bartenschlager (Heidelberg University, Germany) (43). Expression plasmids for DENV2 strain 16681 NS2B3 and NS3, i.e. pcDNA-DENV2-NS2B/3-hco-2xstrep and pcDNA-DENV2-NS3-hco-2xstrep, were gifts from Priya Shah (University of California Davis, USA) (62). Plasmids encoding DENV2 NS2B3 and its protease-inactive mutant, i.e. pcDNA3.1-V5-His-TOPO-TA-DENVNS2B/3-WT and -S135A were obtained from Nicholas J. Lennemann (University of Alabama at Birmingham, Birmingham, AL, USA) (61).

### Transfections and transductions

Cells stably expressing TRIM5α-HA, FLAG-TRIM5α, TRIM56-FLAG or TRIM69-HA were generated by retroviral or lentiviral transduction. Retroviral and lentiviral vectors expressing the TRIMs were prepared by co-transfection in HEK293T cells in 10-cm dishes at 70-80% confluency, using 10 μg of the expression plasmid (pLVX-huTRIM5α-HA, pMIH-FLAG-huTRIM5α, pCX4bsr-TRIM56-WT-FLAG, pCX4bsr-TRIM56-CC21/24AA-FLAG or pMIP-TRIM69-HA) together with 5 μg of pMD2.G (71), and 10 μg of pCl-Eco (72) for retroviral vectors or psPax2 (44) for lentiviral vectors. Transfections were done using polyethylenimine (PEI; Polyscience, Warrington, PA, USA), as described earlier (73). The transfection medium was replaced after 24 h. Viral supernatants were harvested 48 h post-transfection, clarified by low-speed centrifugation, and filtered through a 0.45 μm membrane (Millipore Sigma, Burlington, MA, USA) as described previously (51, 73). Clarified viral supernatants were used to transduce Vero E6, HeLa and HEK293T cells. For this, cells were seeded in 6-well plates at 60% confluency and transduced with 1 ml of each undiluted vector supernatant for 24 h at 37°C. One day later, transduced cells were selected with puromycin (6.5 μg/ml for Vero E6 and 1.5 μg/ml for HeLa and HEK293T; InvivoGen, San Diego, CA, USA) for 3 days (pLVX-and pMIP-derived constructs), hygromycin B (400 μg/ml for Vero E6 and HeLa and 200 μg/ml for HEK293T; Enzo Life Sciences, Farmingdale, NY, USA,) for 7 days (pMIH-derived construct), or blasticidin S (7.5 μg/ml for Vero E6, 5 μg/ml for HeLa and 10 μg/ml for HEK293T; Thermo Scientific, Rockford, IL, USA) for 4 days (pCX4bsr-derived constructs). Untransduced control cells were killed when subjected to these antibiotic treatments.

### DENV production and titration

pFK-DVs-G2A harboring the full-length DENV2 genomic cDNA with the GFP coding sequence inserted 5’ of C (43, 74) was grown in E. coli TOP10 and purified using the PureLink™ HiPure Plasmid Maxiprep Kit (Invitrogen, Carlsbad, CA, USA) as per the manufacturer’s specifications. Restriction digestion using XbaI and ScaI enzymes was used to confirm the integrity of the viral genomic insert. Following linearization of pFK-DVs-G2A as described previously (75), *in vitro* transcription was performed using the mMESSAGE mMACHINE SP6 kit (Thermo Fisher Scientific). Infectious RNA transcripts were electroporated into Vero E6 cells, and supernatants were harvested from 5 to 7 days post-electroporation, filtered (0.45 μm), aliquoted, and stored at −80°C until use. GFP expression, used as a marker of DENV2 infection, was visualized using an Axio Observer microscope (Carl Zeiss, Inc., Toronto, ON, Canada). The titers were expressed as FACS infectious units/mL in Vero E6 cells.

WT DENV2 virus (DENV-WT) strain 16681 was propagated in Vero E6 cells and aliquoted and stored at −80°C, and viral titers were determined by plaque assay as detailed in (76). For this, 24-well plates were seeded with 2 x 10^5^ Vero E6 cells/well. Cells were infected with serially diluted virus samples. Two hours post-infection, inoculums were replaced with DMEM containing 2% carboxymethylcellulose (MP Biomedicals). 7 days after infection, cells were fixed with 10% formaldehyde, and plaques were counted following staining with 1% crystal violet/10% ethanol (Thermo Scientific). Infectious titers are expressed as plaque-forming units per milliliter (PFU/ml).

### Viral challenges and flow cytometry

For DENV2-GFP assays, we first performed infections using serially diluted virus stock (1:1 to 1:64). The amount of virus that yielded 10-20% infection in control untransduced cells was used in subsequent infection assays. Cells were infected for 48 h or 72 h incubation in DMEM medium containing 10% FBS at 37°C at 5% CO2. Following infections, cells were fixed in 4% formaldehyde (Thermo Fisher Scientific) and processed for flow cytometry using a CytoFLEX S flow cytometer (Beckman Coulter Life Sciences, Indiana, USA). Infection was assessed as the percentage of GFP-positive cells. The flow cytometry data were analyzed using FlowJo software version 10.10.0 (BD, FlowJo LLC, Ashland, OR, USA). All infection experiments were performed in triplicate.

### Antibodies and Immunoblotting

For Western blotting (WB) assays, cells were lysed for 10 min at 4°C in lysis buffer (10 mM Tris-HCl, pH 8.0, 1 mM EDTA, 1% Triton X-100, 0.1% Sodium Deoxycholate, 140 mM NaCl) supplemented with Pierce protease inhibitor cocktail (Thermo Fisher Scientific). Boiled lysate samples were subjected to SDS-polyacrylamide gel electrophoresis. Following transfer to nitrocellulose membranes (Bio-Rad Laboratories, Hercules, CA, USA), proteins were probed with rabbit monoclonal anti-FLAG (1:2000, 2368S, Cell Signaling Technology, Danvers, MA, USA), rabbit monoclonal anti-HA (1:5000, 3724S, Cell Signaling Technology), mouse monoclonal anti-DENV NS5 (1:5000, GTX629447, GeneTex, Irvine, CA, USA), rabbit polyclonal anti-DENV NS3 (1:2000, GTX124252, GeneTex), or rat anti-β-actin (1:100000, 664803, BioLegend, San Diego, CA, USA), and revealed using HRP conjugated goat anti-rabbit IgG or horse anti-mouse IgG secondary antibodies (1:5000, 7074S and 7076S Cell Signaling Technology).

### CRISPR/Cas9 knockout

TRIM5, TRIM56, and TRIM69 knockout Huh7 and HeLa cells were generated using the lentiCRISPRv2 (pLCv2) system (69), in which the gRNA, Cas9 nuclease, and an antibiotic resistance marker are co-expressed from a single vector. The previously described pLCv2 plasmid expressing T5gRNA2 was used for TRIM5 knockout (44). lentiCRISPRv2-blast and lentiCRISPRv2-hygro (70) were used for TRIM56 and TRIM69 knockout, respectively. gRNAs targeting exon 3 for TRIM56 and exon 2 for TRIM69 (Table S1 and Fig. S4) were designed according to Zhang’s protocol using CHOPCHOPv3 (77). A control gRNA targeting the non-human CAG hybrid promoter (44) was also used. Sense/antisense pairs of gRNA-encoding ODNs were annealed and cloned into pLCv2 cut with BsmBI. LentiCRISPRv2 clones containing guide sequences were sequenced prior to lentiviral production.

Lentiviral particles were produced by co-transfection of pLCv2, pMD.G and pΔR8.9 in HEK293T cells (44). Viral supernatants were collected at 48 h post transfection and used to transduce Huh7 and HeLa cells. Following transduction, cells were selected with puromycin (1 μg/mL for Huh7, 1.5 μg/mL for HeLa; InvivoGen) for 3 days, blasticidin S (5 μg/mL for Huh7 and HeLa; Thermo Fisher Scientific) for 4 days, or hygromycin B (75 μg/mL for Huh7, 400 μg/mL for HeLa; Enzo) for 6 days, respectively. Control untransduced cells were killed under these conditions.

Genomic DNA was extracted from selected cells using the Geneaid gSYNC DNA extraction kit (Geneaid Biotech, New Taipei City, Taiwan). The regions flanking the targeted sites were PCR-amplified using the locus-specific primer pairs (Table S1). For *TRIM69*, both gRNAs were amplified using a single pair of primers (TRIM69_gRNA1-2_Seq_fwd/rev), generating a 475-bp amplicon which comprises the two target sites, positioned 145 bp apart. For *TRIM56*, primers TRIM56_gRNA1_Seq1_fwd/rev and TRIM56_gRNA2_Seq_fwd/rev generated amplicons of 430 bp and 423 bp spanning the gRNA1 and gRNA2 target sites, respectively. The *TRIM5* target region was amplified using the primers previously described for SURVEYOR analysis of the exon 1 gRNA2 site (44), yielding a 684-bp amplicon. In each case, the primers were positioned to leave at least 150 bp of sequence downstream of the predicted cleavage site, essential for reliable trace decomposition assay. Amplicons were purified and Sanger-sequenced using the forward primer for TRIM56 and reverse primer for TRIM5 and TRIM69 as shown in Table S1. Editing efficiency was quantified from the resulting chromatograms using the TIDE web tool (v3.3.0; https://tide.nki.nl) (78), which quantitates percentages of indels by sequencing decomposition, in comparison with the unedited control CAG or untransduced HeLa or Huh7 cells (Fig. S4).

### RNA extraction and RT-qPCR

Total RNA was extracted from HEK293T cells using TRIzol (Life Technologies, Carlsbad, CA, USA) and chloroform (Sigma-Aldrich, St. Louis, MO, USA), with RNase-free glycogen (Life Technologies) added during extraction to improve RNA yield as described in (74). Extracted RNA was treated with the TURBO DNA-free Kit (Invitrogen) to remove contaminating genomic DNA. One-step RT-qPCR was performed using the One Step TB Green® PrimeScriptTM RT-PCR Kit II (Takara Bio, San Jose, CA, USA). Amplification was performed using 10 μM of forward and reverse primers (DV2-16681s-FWD/REV for DENV and GAPDH-FWD/REV for GAPDH; Table S1) and 2 μl template (100 ng) in 20 μl final volume according to the manufacturer’s protocol. Reverse transcription (RT) was done at 42°C for 5 min followed by 95°C for 10 sec. After RT, initial denaturation was done at 95°C, then 40 cycles of amplification were performed as follows: 5 sec at 95°C, 34 sec at 60°C, followed by dissociation at 95°C for 15 sec, 60°C for 1 min and 85°C for 15 sec. Each PCR was performed in triplicate, and the threshold cycle (Ct) was determined using QuantStudio^TM^ Design&Analysis software v1.5.2 (Thermo Fisher Scientific). Relative expression was calculated using the ΔΔCt method with GAPDH for normalization (2– (*C*t (target)-*C*t (*GAPDH*))).

### Data representation and statistical analyses

Densitometry analyses of western blots were done using Image Lab software (BioRad). Data were plotted and analyzed using GraphPad Prism software v9. Statistical significance was determined by one-way ANOVA followed by Dunnett’s or Tukey’s multiple-comparisons test, as applicable. The specific statistical test used, number of independent experiments, and significance thresholds are indicated in the corresponding figure legend.

## ACKNOWLEDGMENTS

We are most grateful to Kui Li (University of Tennessee, Memphis, TN, USA), Paul D. Bieniasz (The Rockefeller University, New York, NY, USA), Nicholas J. Lennemann (University of Alabama at Birmingham, Birmingham, AL, USA), Ralf Bartenschlager (Heidelberg University, Heidelberg, Germany), Joseph Sodroski (Harvard Medical School, Boston, MA, USA), Feng Zhang (Broad Institute of Massachusetts Institute of Technology and Harvard, Cambridge, MA, USA), Priya Shah (University of California Davis, USA) and Brett W. Stringer (Griffith University, Brisbane, Australia) for the kind gift of reagents. We thank Océanne Mérette (Institut National de la Recherche Scientifique, Laval, Canada) for the preparation of WT DENV2. We also thank Hugo Germain, Tagnon Missihoun and Mélodie B. Plourde for sharing and assistance with instruments.

## AUTHOR CONTRIBUTIONS

R.R.P.: Conceptualization, Methodology, Formal Analysis, Investigation, Data Curation, Writing - Original Draft, Writing - Review & Editing, Visualization

L.L.B.M.: Methodology, Formal Analysis, Investigation

N.M.: Supervision, Methodology, Formal Analysis, Investigation

S.R.: Investigation

A.A.S.: Investigation, Resources

N.H.N.: Formal Analysis, Investigation, Visualization

J.S.: Formal Analysis, Investigation, Visualization

L.C.-C.: Methodology, Resources, Supervision

I.D.-P.: Supervision, Resources

L.B.: Conceptualization, Methodology, Formal Analysis, Resources, Data Curation, Writing - Original Draft, Writing - Review & Editing, Supervision, Project Administration, Funding Acquisition

